# Three-dimensional Multicolor Subcellular Imaging by Fast Serial Sectioning Tomography for Centimeter-scale Specimens

**DOI:** 10.1101/2021.11.11.468237

**Authors:** Wentao Yu, Lei Kang, Victor T. C. Tsang, Yan Zhang, Ivy H. M. Wong, Terence T. W. Wong

**Affiliations:** Translational and Advanced Bioimaging Laboratory, Department of Chemical and Biological Engineering, The Hong Kong University of Science and Technology, Kowloon, Hong Kong, China

## Abstract

Rapid multicolor three-dimensional (3D) imaging for centimeter-scale specimens with subcellular resolution remains a challenging but captivating scientific pursuit. Here, we present a fast, automated, cost-effective, and versatile multicolor 3D imaging method with ultraviolet (UV) surface excitation and vibratomy-assisted sectioning, termed translational rapid ultraviolet-excited sectioning tomography (TRUST). TRUST enables exogenous molecular-specific fluorescence and endogenous content-rich autofluorescence imaging simultaneously with the help of a UV light-emitting diode and a color camera. Commonly applied tissue preparation procedures (e.g., staining or clearing) are laborious, time-consuming, and may induce detrimental effects on processed samples. In TRUST, formalin-fixed specimens are stained with real-time double labeling layer by layer along with serial widefield optical illumination with raster scanning and mechanical sectioning to improve the staining speed and reveal rich biological information. All vital organs in mice have been imaged by TRUST to demonstrate its fast, robust, and high-content multicolor 3D imaging ability. Moreover, its potential for developmental biology has also been validated by imaging entire mouse embryos (taking ∼2 days for imaging the embryo at the embryonic day of 15). TRUST offers a way for multicontrast and multicolor whole-organ 3D imaging with high resolution and high speed while relieving researchers from heavy sample preparation workload.

## Introduction

High-resolution three-dimensional (3D) whole-organ imaging with high fidelity has long been a scientific challenge. Mechanical sectioning followed by manual staining is a traditional approach for 3D histological imaging, which has no limitation on sample size^1^. However, hundreds to thousands of slices are needed to be mounted on glass slides, stained with dyes, and then imaged by a conventional bright-field microscope with raster scanning. The whole procedure is highly labor intensive and requires complicated image registration algorithms^2–7^ for 3D reconstruction. Moreover, sectioning before imaging could induce inevitable distortion in the reconstructed 3D image due to slice ruptures, which is impossible to be corrected by image processing approaches, hindering the wide applicability of this traditional method.

Over the past decade, optical microscopy assisted with mechanical serial sectioning or tissue clearing has been proven as an automated and registration-free solution for large centimeter-scale 3D imaging^8–13^. However, both methods require further development before a translational and rapid multicolor 3D whole-organ imaging system can be implemented. First, simultaneous multicolor imaging of different tissue components can encounter various constraints. For instance, in fluorescence microscopy, it is difficult and time-consuming to achieve a consistent staining performance in both the central and peripheral areas for large and dense tissue samples due to the nature of passive diffusion^14^. Although transgenic fluorescent labeling^15–17^ can solve this issue, it does not apply to normal organs. Besides labeling, the overall cost and complexity of the imaging system could be another concern with the increased number of laser sources and detectors for multicolor imaging^18^. Label-free imaging techniques do not suffer from problems associated with tissue staining. For example, photoacoustic microscopy^8^ can directly image cell nuclei of the whole mouse brain with a 266-nm pulsed ultraviolet (UV) laser. However, its imaging speed is relatively low due to the point-scanning imaging mechanism. In addition, multiple laser sources with different wavelengths are needed to probe different biomolecules. Autofluorescence microscopy is another label-free imaging method that has the ability to image different tissue components based on the autofluorescence signals from endogenous fluorophores, including myelinated fibers^19^, blood cells^20^, lipofuscin^21^, etc. However, the imaging specificity and signal-to-noise ratio (SNR) are relatively low.

To achieve rapid and high-content 3D volumetric imaging, developing advanced and high-speed imaging systems^22–24^ is vitally important. Wide-field imaging techniques, like microscopy with ultraviolet surface excitation (MUSE)^25,26^ or light-sheet fluorescence microscopy (LSFM)^27,28^, generally show faster imaging speed when compared with point-scanning-based approaches, like confocal microscopy^29^ or two-photon microscopy^13^. In addition, eliminating the need for lengthy and laborious sample preparation protocols is crucial as staining or clearing for large specimens, e.g., a whole mouse brain, can take several weeks^7–9^. More importantly, some protocols can induce detrimental effects on processed samples and finally degrade the imaging quality^11,12,14,32–36^. For example, embedding samples in resin or paraffin block^11,12,36^ can result in obvious tissue shrinkage due to dehydration, increasing the difficulty for follow-up tissue staining^12^. Optical tissue clearing also suffers from issues including morphological distortion of the sample^33^, toxicity of reagents^34^, bleaching of fluorophores^34^, and potential loss of some components^35^, although some efforts have been made to reduce the undesired effects^37–39^.

To overcome the aforementioned technical challenges, by building upon MUSE^25,26,40^, here we present a rapid and multicolor 3D microscopy for whole-organ imaging with a light-emitting diode (LED), termed translational rapid ultraviolet-excited sectioning tomography (TRUST). The multicolor and high-content imaging ability of TRUST comes from two parts. First, conventional fluorescent dyes with different emission spectra can be excited simultaneously with a single low-cost UV-LED, providing a broad and informative color palette. In TRUST, two fluorescent dyes (4’,6-diamidino-2-phenylindole (DAPI) and propidium iodide (PI)) have been used together for double labeling^41–43^, which is helpful to provide high color contrast, hence revealing rich biological information (Supplementary Fig. 1). Second, TRUST also enables the imaging of content-rich endogenous fluorophores based on autofluorescence signals which are also excited by the UV-LED, such as myelinated fiber bundles (Fig. 2j) or neurofilaments (Fig. 2p). Collectively, with the help of a color camera, both molecular-specific fluorescence and autofluorescence signals can be captured simultaneously to achieve multicolor and multicontrast imaging.

To avoid the lengthy staining procedure and achieve better staining uniformity, in TRUST, a formalin-fixed sample without any labeling can be directly submerged into the staining solutions in a water tank of a vibratome. Each layer will be labeled followed by serial widefield imaging with raster scanning and vibratomy-assisted sectioning. Considering the non-linear relationship between labeling time and diffusion depth of probes^44–46^, both the speed and uniformity of staining can be significantly improved, especially for large samples.

Our proposed TRUST system can realize fast and high-content multicolor imaging cost-effectively while relieving heavy preparation work and associated undesired tissue changes. To characterize the performance and examine the robustness of the proposed system, several vital mouse whole organs have been imaged. To further show the full potential of TRUST, whole mouse embryos at different stages have also been imaged. The detailed image comparison demonstrates that TRUST is highly desirable for large sample imaging, showing great promise as a tool for developmental biology studies.

## Results

### TRUST system setup and workflow

In TRUST (Fig. 1a–c), the short penetration depth in tissue of obliquely illuminated light from UV-LED (∼285 nm) is the key to achieve wide-field block-face imaging. Both fluorescence and autofluorescence signals from the tissue surface are excited by UV light, which are subsequently collected by a 10× infinity-corrected objective lens, and finally focused on a color camera by a tube lens. Two motorized stages (Motor-*x*, Motor-*y*) can drive the objective lens scanning along the *x-y* plane, and the tunable *z*-axis stage is used for focusing. A lab-built 2-axis angle adjustable platform (Supplementary Fig. 2) under the vibratome can keep the focal plane of the objective lens in parallel with the sample surface determined by the blade angle of the vibratome.

**Figure 1.**
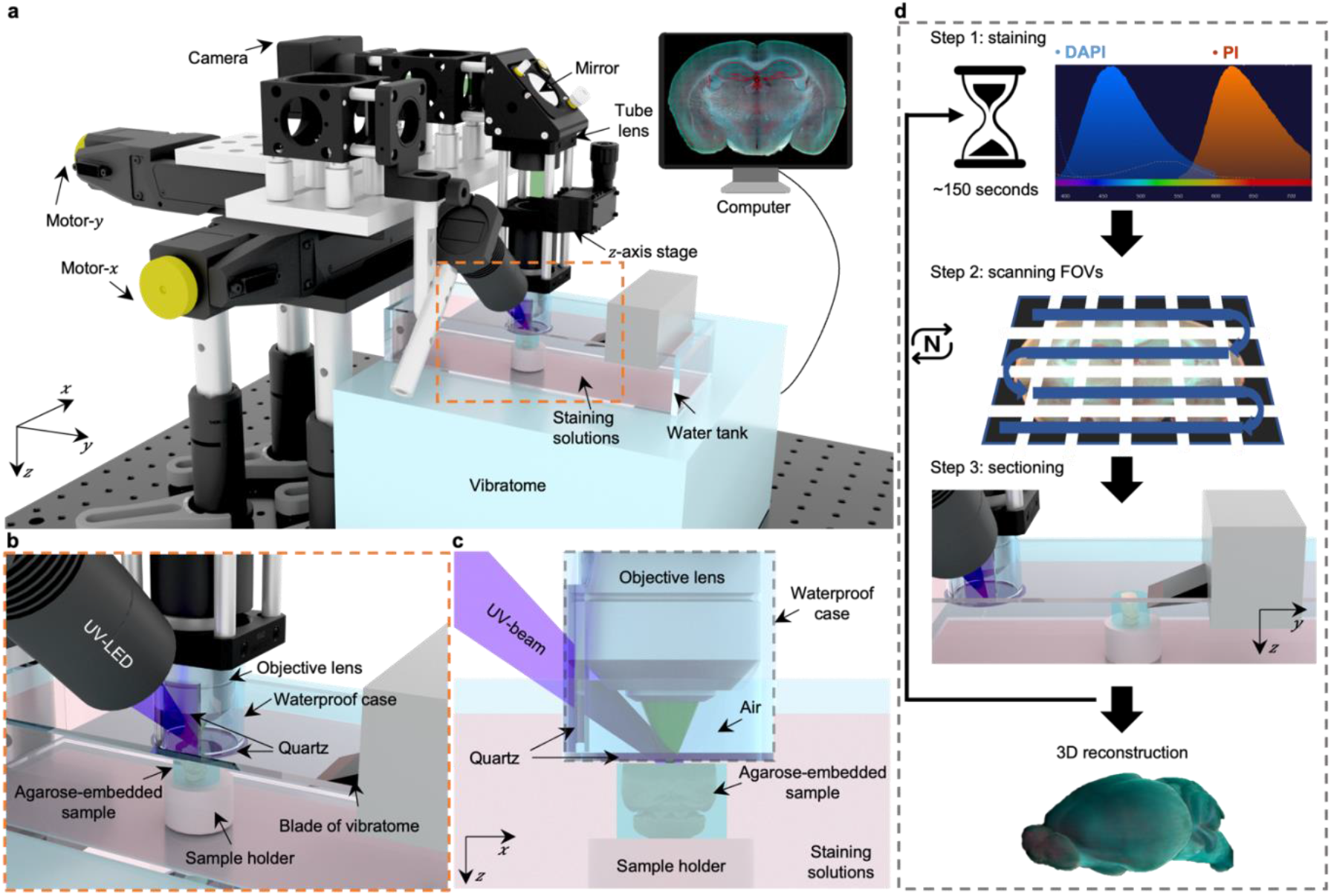
Overview of 3D whole-organ imaging by TRUST. **a**, Schematic of the TRUST system. Light from UV-LED is obliquely projected onto the surface of an agarose-/gelatin-embedded sample (e.g., a mouse brain), which is placed on top of a sample holder inside a water tank of a vibratome filled with staining solutions. The generated fluorescence and autofluorescence signals are collected by an objective lens, refocused by an infinitely corrected tube lens, and finally detected by a color complementary metal-oxide semiconductor camera. **b**, Close-up of the region marked by orange dashed box in a, showing the components that are immersed in the staining solutions. **c**, Viewing b from the *x*-*z* plane. A waterproof case containing two pieces of quartz keeps most of the space under the objective lens filled with air. **d**, Workflow of the whole imaging process, including (1) chemical staining for ∼150 seconds, (2) widefield imaging with raster-scanning and stitching in parallel, and (3) shaving off the imaged layer with the vibratome to expose a layer underneath. The three steps will be repeated until the entire organ has been imaged. All procedures are automated with lab-built hardware and control programs.

Different from conventional fluorescence microscopy, fixed tissue block after agarose or gelatin embedding can be directly imaged with TRUST, and the sample will be stained during imaging by submerging it under staining solutions in the water tank of the vibratome. A 3D printed plastic waterproof case (Fig. 1c) can reduce the fluorescence background by minimizing the amount of staining solutions filled between the tissue surface and objective lens (Supplementary Fig. 3). Besides, it can prevent the objective lens to be affected by liquid evaporation or fluctuation, ensuring high-quality imaging throughout the entire whole-organ imaging process. Two pieces of quartz mounted on the waterproof case are used for the transmission of the UV illumination beam and the excited fluorescence signal.

Workflow of the whole system (Fig. 1d) can be simplified as a loop of three steps: (1) surface layer of the sample immersed under chemical dyes will be labeled for ∼150 seconds (Supplementary Fig. 4); (2) the region of interest of the current layer, which consists of multiple field-of-views (FOVs), will be acquired through motorized raster-scanning and subsequently stitched in parallel by our program; (3) the vibratome will cut off the imaged layer and expose the layer underneath to the staining solutions. This loop will end when the whole organ has been imaged completely. To realize fully automated serial imaging, triggering circuits and corresponding control programs have been developed for synchronizing the entire TRUST system.

Because the sample is stained during the imaging step in TRUST, the labeling protocol can be thought of as real-time staining. Two chemical dyes, DAPI and PI were used together to stain nucleic acid in cells. The fluorescence intensity of DAPI or PI will be amplified over 20 folds when they are bonded with the nucleic acids^47,48^, so that the fluorescence background from the staining solutions will be negligible. Finally, to maintain the concentration of chemical dyes in the water tank over a long period of time for whole-organ imaging, a high-precision water pump (LabN1-YZ1515x, Baoding Shenchen Precision Pump Co., Ltd) can be utilized to add additional dyes into the water tank at a certain rate (Supplementary Table 1). Another peristaltic pump (KCP PRO2-N16, Kamoer Fluid Tech Co., Ltd.) can be utilized to automatically collect the sectioned layers for follow-up studies if necessary (Supplementary Movie 1).

### Whole mouse brain imaging with TRUST

A mouse brain was first imaged by our TRUST system to demonstrate the fast imaging speed, molecular-specific real-time staining, and the multicolor imaging capability. The mouse brain was first harvested and fixed in formalin for 24 hours. Subsequently, the mouse brain was embedded into 2% w/v agarose. The sectioning thickness of the vibratome was set as 50 µm to provide a good balance between the sectioning quality and the *z*-axis sampling interval. DAPI and PI solutions with a concentration of 5 µg/ml were used to label cell nuclei. The total imaging volume (12.1 mm × 8.6 mm × 17.4 mm (*xyz*)) consists of approximately 7.8 × 10^11^ voxels with 24 bits RGB channels, of which the uncompressed dataset is ∼2.1 terabytes (TB). Without the need for image registration, images of 347 coronal sections acquired by TRUST can be directly stacked to reconstruct the 3D model (Fig. 2a). Including staining, two-dimensional (2D) raster-scanning, and mechanical sectioning, the total acquisition time is ∼64 hours, which is highly manageable and practical. In comparison, the whole-brain staining or clearing in conventional fluorescence imaging will already take weeks^9,30,49^, not to mention the follow-up optical scanning.

**Figure 2.**
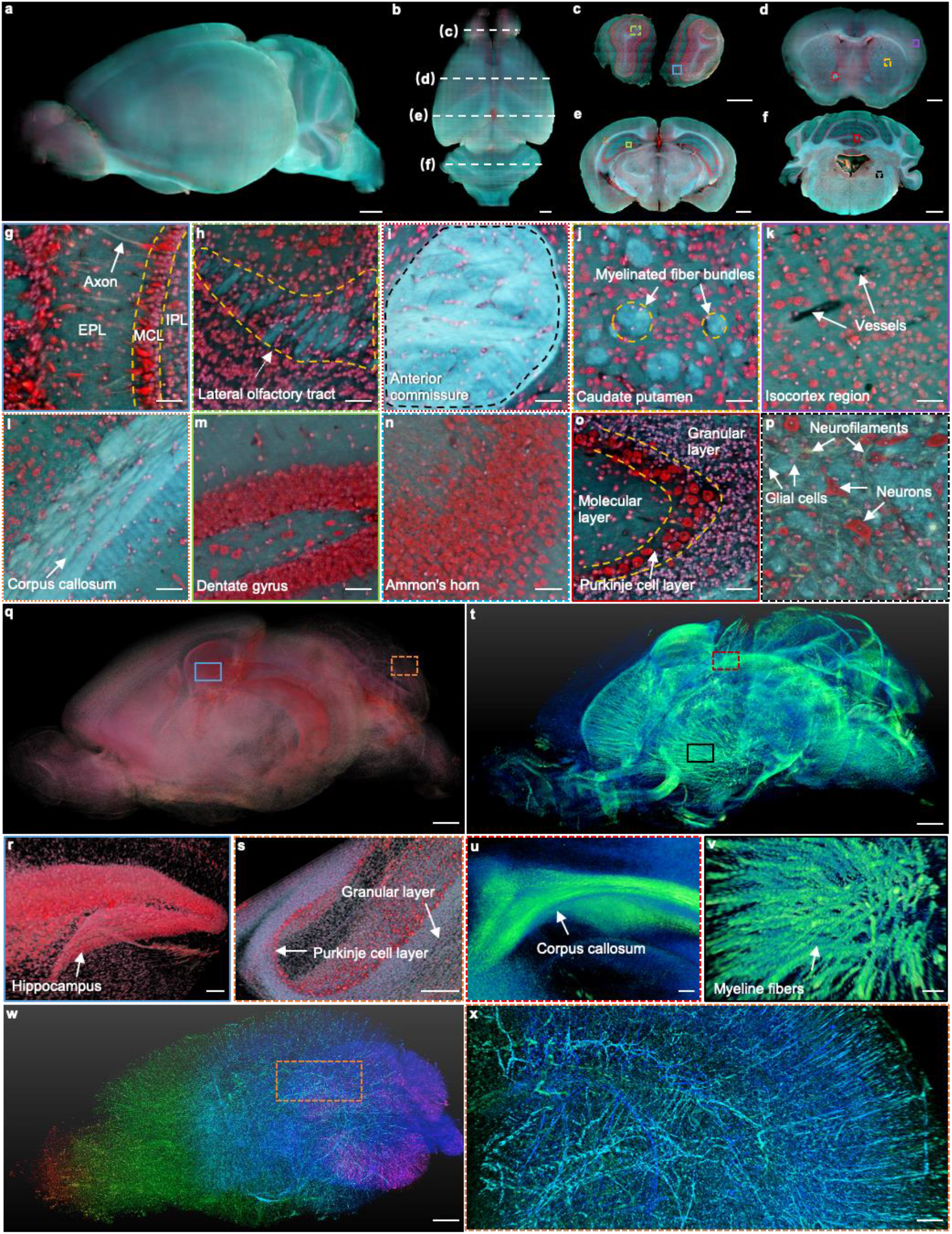
Whole mouse brain imaging with TRUST. **a, b**, Side and top views of the reconstructed 3D model of a fixed mouse brain, respectively. **c–f**, Four coronal sections with positions indicated in b by the white dashed lines. **g, h**, Close-up images of the blue solid and green dashed regions marked in c. **i–k**, Close-up images of the red dotted, yellow dashed, and purple solid regions marked in d. **l–n**, Close-up images of the orange dotted, green solid, and blue dashed regions marked in e. **o, p**, Close-up images of the red solid and black dashed regions marked in f. **q**, The cell nuclei in the mouse brain extracted from the red channel in a. **r, s**, Close-up images of the blue solid and orange dashed regions marked in q. **t**, 3D structure of the nerve tracts or fibers extracted from the green channel of the autofluorescence signal in a. **u**,**v**, Close-up images of the red dashed and black solid regions marked in t. **w**, Vessel network extracted from another mouse brain and rendered with different colors corresponding to different coronal layers. **x**, Close-up image of the orange dashed region marked in w. MCL: mitral cell layer; IPL: inner plexiform layer; EPL: external plexiform layer. Scale bars: 1 mm (**a–f, q**,**t**,**w**), 200 µm (**x**), 100 µm (**r–v**), and 50 µm (**g–p**).

To compare the imaging speed of TRUST with other systems (Supplementary Table 2), sample preparation time has also been counted because it generally takes much longer time than that of optical scanning, especially for centimeter-sized samples (e.g., a whole mouse brain). To this end, the imaging speed of TRUST for the whole mouse brain can be up to ∼0.8 TB/day, which is faster than most of the previously reported high-resolution 3D imaging systems, including serial two-photon tomography^50^ (∼0.2 TB/day), microtomy-assisted photoacoustic microscopy^8^ (∼0.002 TB/day), micro-optical sectioning tomography^11^ (∼0.2 TB/day), fluorescence micro-optical sectioning tomography^36^ (∼0.3 TB/day), and LSFM with CUBIC^51^ (∼0.5 TB/day).

Four coronal sections (Fig. 2c–f) with positions indicated by the white dashed lines in Fig. 2b show the stable sectioning performance of vibratome and uniform staining throughout the whole-brain imaging process. With traditional staining protocols^9,46^, the center area is harder to be labeled compared with its peripheral area (Supplementary Fig. 5a) due to the nature of passive diffusion of stains. Besides, agents like Triton X-100, which is used for increasing the permeability of tissue and accelerating the staining speed, can also increase the transparency of the sample by washing away the lipids inside. These protocols are not suitable for UV surface excitation-based imaging systems because the imaging contrast can then be deteriorated (Supplementary Fig. 5b–g).

In addition to the high imaging speed, the advantages of TRUST are more reflected in its high-content imaging ability with a low cost by taking the advantages of the UV surface excitation and double labeling (Supplementary Fig. 1 and supplementary Table 2). Conventionally, multiple lasers with several sets of excitation/emission filters are needed in fluorescence microscopy to achieve multicolor imaging^46^. In the case of the TRUST system, a single UV-LED and a typical color camera are used for the excitation and detection, respectively, because most of the conventional fluorescent dyes can be excited by deep UV light and emit photons in the visible range^25^. Different from the staining protocol used in MUSE for rapid 2D slide-free histology^25^, we developed a real-time double-labeling protocol to make the staining perfectly fitted with our TRUST system. The double labeling with DAPI and PI helps reveal more biological information, achieving better image quality. First, PI staining can clearly reveal cytomorphological details of neurons while the cell bodies of glial cells are only slightly labeled^52^, hence differentiating neurons from glial cells in the brain through morphological differences (Supplementary Fig. 1a). In contrast, DAPI almost only stains cell nuclei. As a result, with the combination of DAPI and PI, TRUST can provide similar imaging contrast as Nissl stains (Supplementary Fig. 6), where the Purkinje cell layer can be differentiated from the granular layer or molecular layer. Furthermore, the color contrast of images can be improved by double labeling, especially for regions with a high density of cells (Supplementary Fig. 1g,h) because the cytoplasm stained by PI (red) can act as a background for DAPI-labeled cell nuclei to stand out (green and blue).

Autofluorescence signals from endogenous biomolecules have shown great promise in revealing biological information in 2D label-free imaging^19–21,53,54^, which have also been utilized in TRUST. For example, the axon (Fig. 2g), nerve tracts (e.g., lateral olfactory tract (Fig. 2h) or anterior commissure (Fig. 2i)), and myelinated fiber bundles in the caudate putamen (Fig. 2j) have been identified with TRUST even without any labeling. Also, blood vessels (Fig. 2k) can be identified based on negative contrast because its autofluorescence signal is lower than that of the surrounding tissues^55–57^. Although autofluorescence signals can be treated as background noise which can deteriorate the fluorescence imaging quality, in TRUST, due to the usage of the double labeling and color camera, the mixed dyes with a broad emission spectrum (blue to red) make the fluorescence signal less affected by the autofluorescence background by extracting signals from different color channels for different types of tissue components. For example, although hepatocytes in liver tissue are hard to be differentiated from autofluorescence background in the red channel, the image contrast is high in the green and blue channels (Supplementary Fig. 7a–d). Also, although the autofluorescence background is strong in the green and blue channels for some regions in the mouse brain, the cells remain to be clear in the red channel (Supplementary Fig. 7e–h).

With background subtraction and dynamic range adjustment in different color channels of TRUST images, the 3D distribution of cell nuclei (Fig. 2q–s) or nerve fiber bundles (Fig. 2t–v) in the brain can be digitally extracted. To further enhance the contrast of the vessel network in the brain (Fig. 2w,x), the mouse was perfused transcardially with a mixture of black ink and 3% w/v gelatin (Supplementary Fig. 8). The 3D animation of the whole mouse brain, including serial coronal or transverse sections, has been rendered as shown in Supplementary Movie 2. The vessel network has also been rendered as shown in Supplementary Movie 3.

### 3D imaging of other organs with TRUST

To demonstrate the generalizability and robustness of the TRUST system, other mouse organs with various sizes are imaged, including heart (Fig. 3a–e), liver (Fig. 3f–j), kidney (Fig. 3k–o), lung (Fig. 3p– t), and spleen (Fig. 3u–y). A fixed mouse heart was first imaged by TRUST with a sectioning thickness of 50 µm. The entire imaging and staining automated procedure for the entire volume (10.4 mm × 8.2 mm × 6.1 mm, 1.8×10^9^ voxels with 24 bits RGB channels, 122 sections) took ∼21 hours, and the reconstructed 3D model is shown in Fig. 3a. One representative section is shown in Fig. 3b with its position marked by a white dashed line at the top right-hand corner. Three close-up images (Fig. 3c–e) indicate that not only the cell nucleus, other components, like adipose tissue (Fig. 3c) or cardiac muscle (Fig. 3d), can also be clearly imaged. More details of the whole dataset, including multiple zoomed-in regions, can be found in the rendered 3D animation (Supplementary Movie 4).

**Figure 3.**
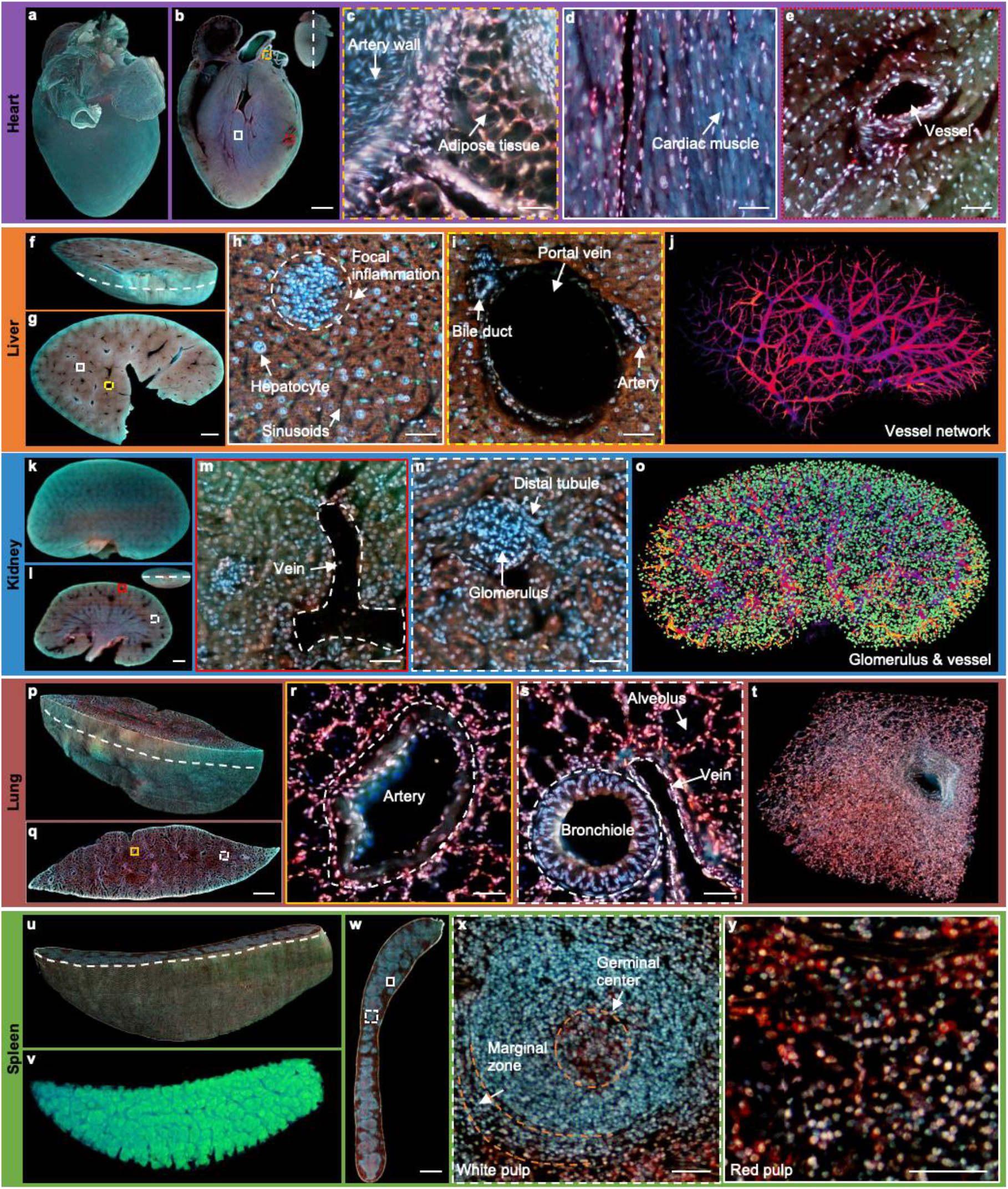
2D/3D image gallery of other organs in mice with TRUST. **a**, Reconstructed 3D model of the whole mouse heart. **b**, One section of the heart with the position indicated by the white dashed line at the top right-hand corner. **c–e**, Three close-up images of the orange dashed, white solid, and red dotted regions marked in b. **f**, One block of a fixed mouse liver. **g**, One section of the liver with the position indicated by the curved white dashed line marked in f. **h**,**i**, Close-up images of the white solid and yellow dashed regions marked in g. **j**, Vessel network extracted based on negative contrast and rendered with pseudo color. **k**, Reconstructed 3D model of a whole mouse kidney. **l**, One section in the middle of the kidney with the position indicated by the white dashed line at the top right-hand corner. **m**,**n**, Close-up images of the red solid and white dashed regions marked in l. **o**, Vessel network and glomeruli extracted from the whole kidney and rendered with pseudo color. **p**, Part of a mouse lung imaged with TRUST. The curved white dashed line indicates the position of the section in q. **r**,**s**, Close-up images of the orange solid and white dashed regions marked in q. **t**, A small zoomed-in region in p. **u**, Rendered 3D model of a mouse spleen and the curved white dashed line indicates the position of the section in w. **v**, White pulps extracted from u through the blue channel and rendered with pseudo color. **x**,**y**, Close-up images of the white dashed and white solid regions marked in w. Scale bars: 1 mm (**b**,**g**,**l**,**q**,**w**) and 50 µm (**c–e**,**h**,**i**,**m**,**n**,**r**,**s**,**x**,**y**).

Part of a fixed mouse liver (Fig. 3f) has also been imaged by TRUST with a sectioning thickness of 50 µm. The entire imaging and staining time for the total volume (8.8 mm × 12.5 mm × 3 mm, 1.4×10^11^ voxels with 24 bits RGB channels, 60 sections) took ∼11 hours. The curved white dashed line marked in Fig. 3f indicates the position of the section in Fig. 3g. Two close-up images (Fig. 3h,i) of the white solid and yellow dashed regions in Fig. 3g show that hepatocytes and typical anatomical structures, like portal vein, sinusoids, and focal inflammation, can be well differentiated with TRUST. Based on the negative contrast of blood vessels as shown in Fig. 3i, the vessel network (Fig. 3j) of the whole dataset can also be extracted with several image processing steps (Supplementary Fig. 9). 3D animation of the whole dataset has been rendered as shown in Supplementary Movie 5. Moreover, another block of the liver with a volume of 500 µm × 500 µm × 250 µm (Supplementary Fig. 10) has also been imaged by TRUST with a sectioning thickness of 10 µm. The fine sectioning thickness results in a decrease in the *z*-sampling interval, which is helpful for resolving microstructures, such as the sinusoids in 3D space.

Imaging for the whole mouse kidney (Fig. 3k) took ∼19 hours and the sectioning thickness of the volume (13.2 mm × 8.6 mm × 4.7mm, 2.3×10^11^ voxels with 24 bits RGB channels, 93 sections) was also set as 50 µm. One typical section is shown in Fig. 3l with its position marked by the white dashed line at the top right-hand corner. Vein and glomerulus can be clearly recognized as shown in Fig. 3m,n, respectively. All glomeruli in the kidney were extracted with the assistance of an open-source object detection library, Detectron2^58^. The reconstructed vessel network together with the glomeruli of the whole kidney is shown in Fig. 3o. Video of the whole dataset has been rendered as shown in Supplementary Movie 6.

Sectioning lung tissue directly with a vibratome can be difficult due to the presence of porous structures, e.g., alveoli or bronchioles (Supplementary Fig. 11a,b). Lung inflation^59–61^ with hydrogels (e.g., agarose or gelatin) injection through the trachea is a common solution to preserve its morphology, thus achieving better sectioning performance. In TRUST, we first filled the lung with 10% w/v melted gelatin by syringe injection, and the processed lung sample should be cooled down quickly and moved into formalin solution at 4 °C overnight for post-fixation (Supplementary Fig. 11c). The entire imaging and staining time for the total volume (15.4 mm × 4.7 mm × 4.6 mm, 1.4×10^11^ voxels with 24 bits RGB channels, 91 sections) took ∼11 hours with a sectioning thickness of 50 µm, and its reconstructed 3D model is shown in Fig. 3p. To exhibit the improved sectioning performance, all sections of the whole dataset including multiple zoomed-in regions have been rendered in the 3D animation video (Supplementary Movie 7). Two close-up images (Fig. 3r,s) of the yellow solid and white dashed regions marked in Fig. 3q show that the common anatomical structures, including artery, bronchiole, and alveolus, can be well imaged.

The final whole organ that we imaged was a mouse spleen. The entire imaging and staining time for the total volume (12.7 mm × 8.2 mm × 3.8 mm, 1.7×10^11^ voxels with 24 bits RGB channels, 75 sections) of the mouse spleen embedded in gelatin block took ∼14 hours. The reconstructed 3D model is shown in Fig. 3u, and the curved white dashed line indicates the position of the section shown in Fig. 3w. Two close-up images (Fig. 3x,y) of the white dashed and white solid regions marked in Fig. 3w correspond to two major components in the spleen: white pulp and red pulp, respectively. Typical structures, like germinal center or marginal zone, can be clearly differentiated. As the overall appearance of the white pulp region (Fig. 3x) looks blue, all white pulp of the dataset as shown in Fig. 3v can be extracted from the blue channel with simple thresholding. Finally, 3D animation of the whole dataset also has been rendered as shown in Supplementary Movie 8.

### 3D imaging of whole mouse embryos with TRUST

Under embryonic development, there are dramatic changes in cell/structural morphology and arrangement. Therefore, 3D imaging of a whole mouse embryo is vitally important to assist biologists to have a complete understanding of any anatomical and functional changes. However, imaging a large size embryo is a scientific challenge. Although effort has been spent by many research groups, their approaches still suffer from either (1) low imaging resolution (micro-CT^62^ and optical projection tomography^63^), (2) a limited number of sections (histological imaging^64–66^), (3) limited sample size due to time-consuming clearing and/or staining (selective plane illumination microscopy^67^ and confocal microscopy^68^). In the above, we have already demonstrated the advantages of our TRUST system, especially its ability for handling various types of organs regardless of their size, and its high imaging speed due to the real-time staining and wide-field imaging configuration. Therefore, the TRUST system theoretically has the potential for addressing the needs in developmental biology.

We first imaged two heads from two mouse embryos with intact skulls (embryonic day (E) 15: Fig. 4a–f; E18: Fig. 4g– l). The whole imaging procedure (including staining, scanning, and sectioning) for the embryo head at E15 (9.9 mm × 9 mm × 6.9 mm, 2.6×10^11^ voxels with 24 bits RGB channels, 137 sections) took ∼20 h, and the reconstructed 3D model from different view angles is shown in Fig. 4a. To achieve better sectioning performance with the vibratome, the skin outside the embryo head at E18 was manually removed, and it took ∼32 hours for both imaging and staining (9.4 mm × 7.8 mm × 10 mm, 3.1×10^11^ voxels with 24 bits RGB channels, 201 sections). By comparing the two datasets, it is possible to determine the evolution of main morphological features over different developing stages. For example, close-up images of the retina areas from the two embryos are compared as shown in Fig. 4e (E15) and Fig. 4k (E18). It can be observed that the inner plexiform layer (IPL) becomes obvious only in the E18 embryo which is consistent with the neural differentiation status of the retinal region^69^. Also, Fig. 4f and Fig. 4l correspond to the cerebral cortex regions in the brain of the E15 embryo and E18 embryo, respectively. The ventricular zone (VZ) is easier to be differentiated from the intermediate zone (IZ) in the E18 embryo due to the differentiation of cortical neuroectoderm^70^. Similar results have also been observed in histological images^65^, proving the images from TRUST is highly reliable.

**Figure 4.**
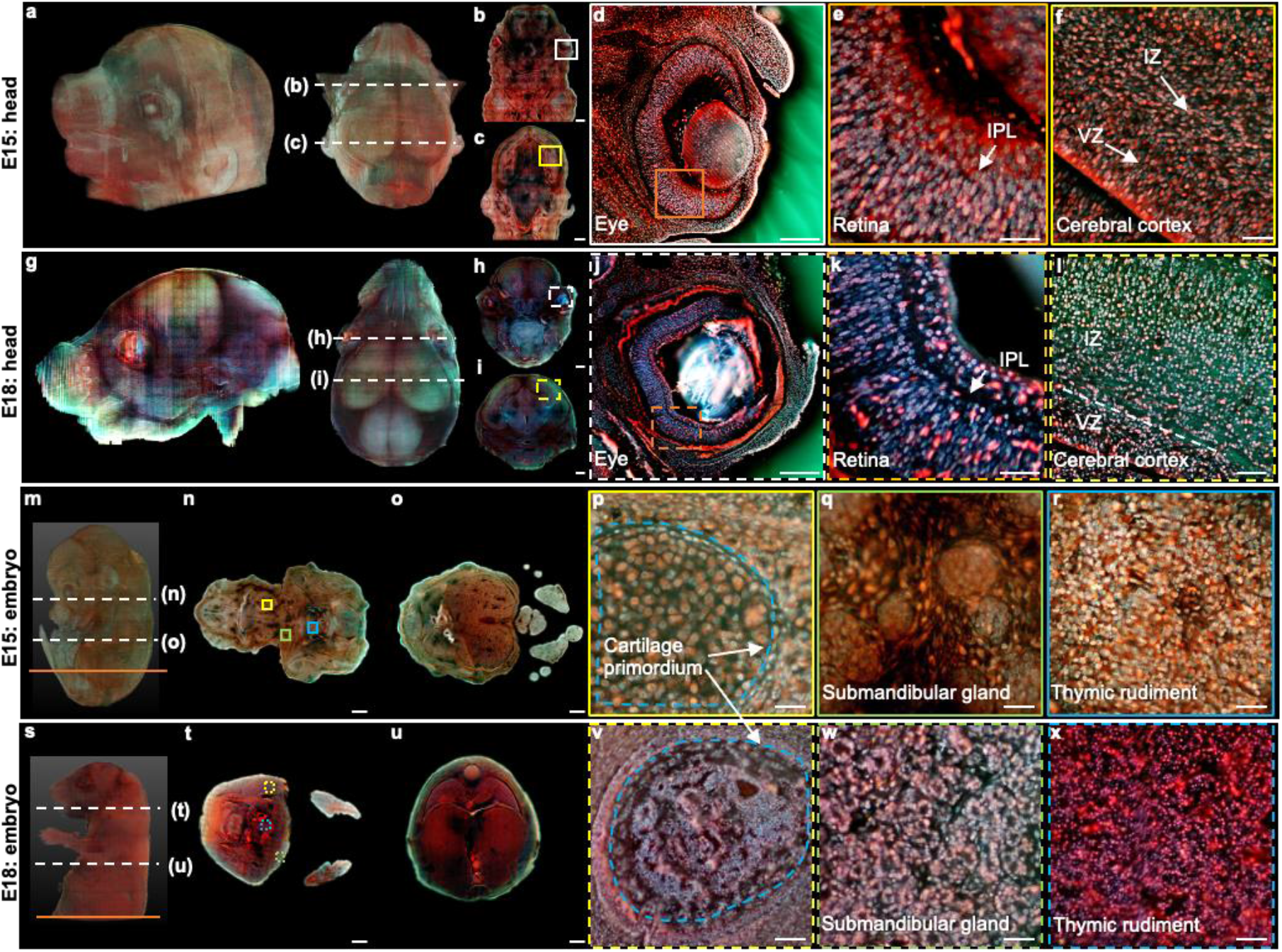
3D imsaging of mouse embryos. **a**, Rendered 3D model of the embryo head at E15 viewed from two perspectives. The two white dashed lines indicate the positions of two sections in b,c. **d**, Close-up image of the white solid region marked in b. **e**, Close-up image of the orange solid region marked in d. **f**, Close-up image of the yellow solid region marked in c. **g**, Rendered 3D model of the embryo head at E18 viewed from two perspectives. The two white dashed lines indicate the positions of two sections in h,i. **j**, Close-up image of the white dashed region marked in h. **k**, Close-up image of the orange dashed region marked in j. **l**, Close-up image of the yellow dashed region marked in i. **m**, Rendered 3D model of a whole mouse embryo at E15. The two white dashed lines indicate the positions of two sections in n,o. **p–r**, Close-up images of the yellow, green, and blue solid regions marked in n. **s**, Rendered 3D model of a whole mouse embryo at E18. The two white dashed lines indicate the positions of two sections in t,u. **v–x**, Close-up images of the yellow, green, and blue dashed regions marked in t. Scale bars: 1 mm (**b**,**c**,**h**,**i**,**n**,**o**,**t**,**u**), 500 µm (**d**,**j**), and 50 µm (**e**,**f**,**k**,**l**,**p–r**,**v–x**).

Then, we have performed 3D imaging of two whole mouse embryos using our TRUST system with a sectioning thickness of 50 µm. The total imaging and staining time for the E15 embryo (9.9 mm × 7.8 mm × 12.8 mm, 4.2×10^11^ voxels with 24 bits RGB channels, 256 sections) took ∼2 days (Fig. 4m–r). The skin of the E18 embryo was also removed, and the entire imaging and staining time (13.2 mm × 9.8 mm × 16.8 mm, 9.3×10^11^ voxels with 24 bits RGB channels, 336 sections) took ∼3 days (Fig. 4s–x). More anatomical differences can be found between the E15 and E18 embryos. For example, close-up images of the cartilage primordium regions from the E15 embryo (Fig. 4p) and E18 embryo (Fig. 4v) indicate that ossification may happen between E15 and E18 stages. Also, the distinguish morphological difference between the submandibular gland in E15 embryo (Fig. 4q) and E18 embryo (Fig. 4w) is consistent with the fact that the acini in a mouse embryo will not complete lumenization until E17^71^. Finally, because of differentiation, cell nuclei in the thymic rudiment regions of the E15 embryo (Fig. 4r) are obviously larger than that in the E18 embryo (Fig. 4x), which is also consistent with the histological images in eHistology Atlas^72^ (Supplementary Fig. 12).

To better illustrate the difference between the two embryos, two whole sections from two embryos with positions marked by the orange solid lines in Fig. 4m and Fig. 4s are shown in Fig. 5. More specifically, Fig. 5a1 and Fig. 5a2 are the zoomed-in regions of the dorsal root ganglion near the spinal cord. The cell nuclei in the latter one look relatively smaller due to cell differentiation. Fig. 5b1 and Fig. 5b2 are the close-up images of the kidney area. Many ureteric buds and almost no glomeruli were observed in the E15 embryo while glomeruli are easily observed in the E18 embryo. The result is consistent with the fact the formation of glomeruli approximately begins at E13.5 with its number subsequently increasing up to 54 folds from E13.5 to E17.5^73^. Fig. 5c1 and Fig. 5c2 are the zoomed-in regions of the spinal cord. The central canal in the former one is larger than that of the latter one, which has also been observed through histological imaging^65^. Fig. 5d1 and Fig. 5d2 are the zoomed-in regions of the adrenal gland. Due to cell differentiation, the cell nuclei in the former one are obviously larger than that of the latter one. Fig. 5e1 and Fig. 5e2 are the zoomed-in regions of the intestines. The muscle structure in the latter one is more obvious than that of the former one. All these high-resolution and rich-molecular contrast TRUST images show that TRUST is an ideal tool for basic biology studies.

**Figure 5.**
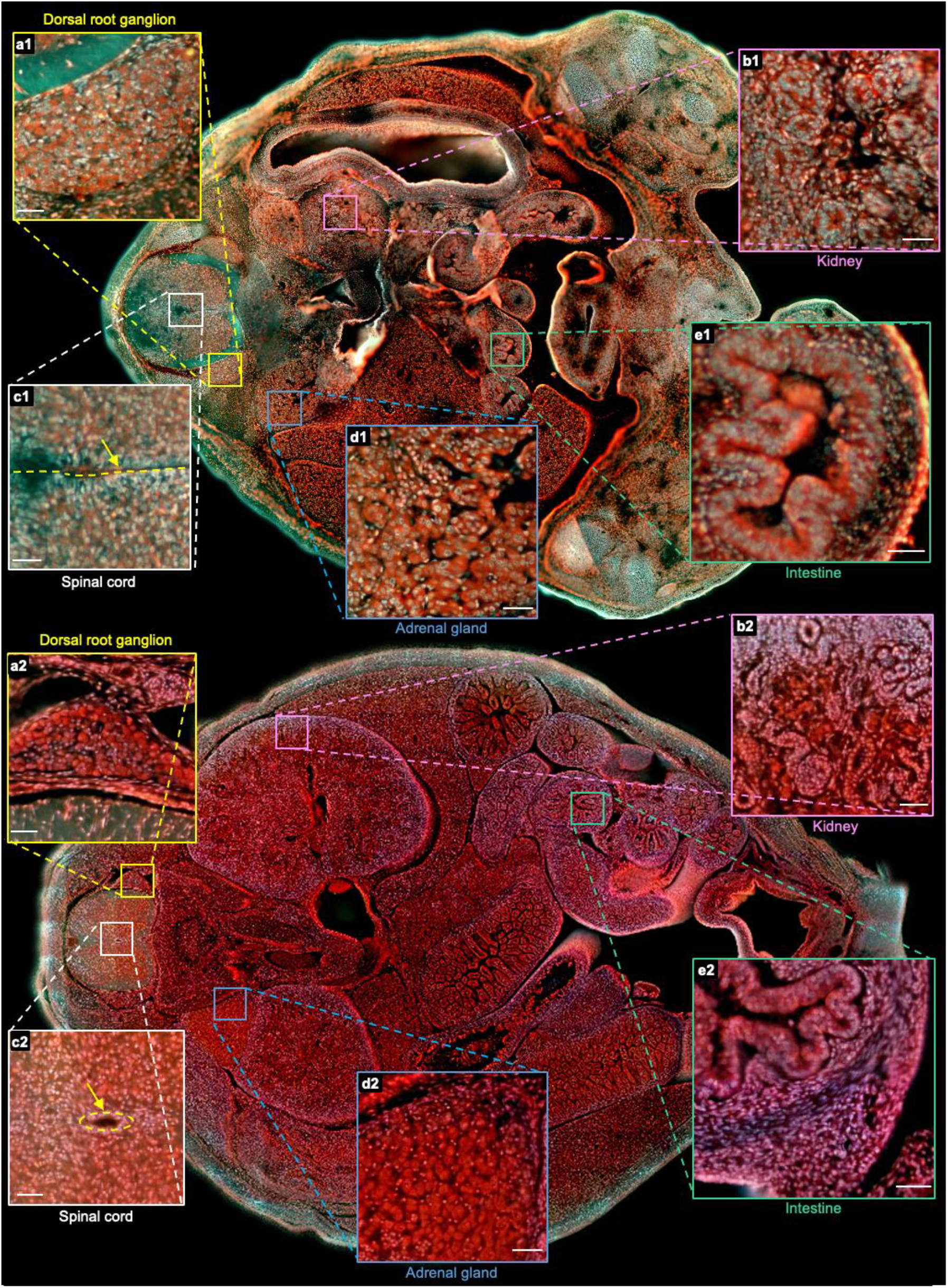
Comparison of two sections from the E15 and E18 embryos. **a1(a2)–e1(e2)**, Close-up images from the E15(E18) embryo of the dorsal root ganglion, kidney, spinal cord, adrenal gland, and intestine marked with yellow, pink, white, blue, and green solid boxes, respectively. Scale bars: 50 µm.

## Discussion

In summary, using our proposed TRUST system, common sample preparation procedures are not required, including dehydration, delipidation, decalcification, or pre-staining. By eliminating the needs of all these time-consuming and labor-intensive sample preparation work, TRUST can provide high-quality subcellular imaging in a time-effective manner. The high-content images acquired by the TRUST system are enabled by the double labeling and UV excited fluorescence and autofluorescence signals. To show the high performance and robustness of the TRUST system, a whole brain from an adult mouse and several other organs (heart, liver, kidney, lung, and spleen) were all imaged by TRUST with high-quality and distortion-free images. Finally, two whole mouse embryos at developing stages of E15 and E18 have also been imaged. The structural and morphological comparison of the two sets of images shows that TRUST is a promising tool for embryonic development study. TRUST is at its early developing stage. To improve the performance of the system, future research development can be carried out in the following aspects:

Firstly, the imaging resolution of TRUST is anisotropic **—** by imaging fluorescent particles with an average diameter of 0.2 µm as shown in Supplementary Fig. 13, the lateral resolution of the TRUST system with a 0.25 numerical aperture objective lens can be roughly estimated by the full width at half maximum of the Gaussian fitted curve, which is ∼1.25 µm. However, the axial resolution of TRUST is several tens of micrometers, which is limited either by (1) the sectioning thickness of the vibratome or (2) the optical-sectioning thickness provided by UV surface excitation. For the sectioning thickness, although a small block of the liver (Supplementary Fig. 10) has been successfully imaged by TRUST with 10-µm interval, which is also the minimal sectioning thickness claimed by the vibratome (VF-700-0Z, Precisionary Instruments Inc.), lots of conditions have to be met (e.g., the sharpness of the blade, the hardness of the tissue and working parameters of vibratome) and its performance is not stable. Therefore, to produce even and consistent sections, the sectioning thickness for other samples is all set as 50 µm. In the future, a more advanced vibratome with better performance (e.g., VF-300-0Z, Precisionary Instruments Inc.) can be used to alleviate this problem. Whereas for the optical-sectioning thickness, it is provided by the UV surface excitation (10∼20 µm^12,25,40,74^), which is relatively large and can vary with different types of tissues. By applying water immersion illuminance, the penetration depth of UV light can achieve ∼50% reduction^40^. Also, doping the sample with UV-absorbent dye can even improve the axial resolution to ∼1 μm^12^. However, these methods also impose stringent requirements for mechanical sectioning. Therefore, integrating optical sectioning methods, like structured illumination^75,76^, could be one of the most promising solutions which can improve the axial resolution and relax the requirement for the mechanical sectioning simultaneously.

Besides, the total scanning time (*T*_0_) for each section of the TRUST system can be further reduced, which can be calculated as:

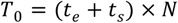

where *t*_*e*_ is the exposure time for each FOV, *t*_*s*_ is the motor scanning time required to move from the first FOV to the next FOV, and *N* is the total number of FOVs. Currently, *t*_*e*_is relatively large (∼250 ms) to maintain an acceptable SNR. Employing multiple light sources (e.g., four UV-LEDs) and illuminating the samples from different angles simultaneously^26^ is a feasible solution to decrease the *t*_*e*_ by a factor of two. Also, *t*_*s*_ can be reduced, at the least, by a factor of two through updating the current motor (20 mm/s; L-509, PI miCos GmbH) to an advanced model (90 mm/s; L-511, PI miCos GmbH). Moreover, the FOV is relatively small (600 mm × 450 mm) with the current camera (DS-Fi3, Nikon Corp.; chip size: 6.91 mm × 4.92 mm) and 10× objective lens. Upgrading the camera with a large chip size (e.g., GS3-U3-123S6C-C, Edmund Optics Inc.; chip size: 14.13 mm × 10.35 mm) and using a low magnification (e.g, 4×) objective lens can reduce the *N* by a factor of ∼25. As for the degraded imaging resolution, it can be later recovered with the help of deep-learning-based super-resolution networks^77–79^. In summary, *T*_0_ can be potentially reduced by a factor of ∼50.

In addition, the imaging contrast can be improved to distinguish more types of components. TRUST images contain both fluorescence and autofluorescence information. To enrich this molecular labeling, any dyes that fluoresce strongly only upon binding to target are potentially compatible with real-time staining. In addition to the cell nucleus, other components in tissue, like protein (8-Anilino-1-naphthalene sulfonic acid, Sigma-Aldrich) and lipids (Nile Red, Sigma-Aldrich) can also be labeled. Furthermore, autofluorescence spectra of some tissue components can overlap with each other^80^, making them hard to be differentiated. Automatically switching between different spectral windows according to the desired imaging target by electrically tunable band-pass filters (e.g., KURIOS-VB1, Thorlabs Inc.) is one possible solution, despite that the exposure time for each FOV could increase because of time multiplexing.

Finally, the staining protocols can be optimized. Choosing dye with high fluorescence enhancement factors upon binding to its target, e.g., SYTOX® Blue (Thermo Fisher Scientific Inc., 500-fold improvement) and YOYO™-1 (Thermo Fisher Scientific Inc., >1000-fold improvement), can further decrease the background signal caused by the staining solutions. Another benefit that comes with it is that the concentration of the staining solutions can be further increased to accelerate the staining speed. Also, the speed control of adding additional dyes with the pump for maintaining the concentration of the stains can be more accurate through negative feedback: the pumping speed will be automatically adjusted with the change of average brightness of each 2D TRUST image.

## Methods

### TRUST imaging system

The short penetration depth of UV light is utilized for 2D block-face imaging. By combining with a vibratome, the TRUST system can realize serial surface imaging followed by sectioning for the whole sample. The schematic of the TRUST system is shown in Fig. 1a–c. Firstly, UV light from a mounted LED (M285L5, Thorlabs Inc.) is slightly focused on the surface of the specimen by a pair of lenses (#67-270, Edmund Optics Inc.; LA4306-UV, Thorlabs Inc.) with an oblique orientation. Then, excited fluorescence and autofluorescence signals will be collected by a 10× objective lens (RMS10X, Olympus Corp.), refocused by an infinity-corrected tube lens (ACA254-200-A, Thorlabs Inc.), reflected by an aluminum mirror (PF10-03-G01, Thorlabs Inc.), and finally detected by a color camera (DS-Fi3, Nikon Inc.). An optical long-pass filter can be placed before the tube lens to further reduce the backscattered UV light. A custom-built 2-axis angle adjustable platform (Supplementary Fig. 2) under the vibratome (VF-700-0Z, Precisionary Instruments Inc.) ensures the focal plane of the objective lens is parallel with the surface of the sample sectioned by the vibratome. Two motorized stages (Motor-*x*: L-509.20SD00, Motor-*y*: L-509.40SD00; PI miCos GmbH) and a manually tunable *z*-axis stage are used for 2D raster-scanning and objective lens focusing, respectively. A 3D printed plastic waterproof case (Fig. 1c) can reduce the fluorescence background from the staining solutions, hence improving the imaging contrast (Supplementary Fig. 3), by keeping most of the space between the objective lens and sample surface filled with air instead of staining solutions. A waterproof case can also keep the imaging plane uninfluenced by the fluid level fluctuations induced by mechanical scanning and liquid evaporation. Two pieces of quartz glasses mounted on the waterproof case are used for the transmission of both UV light and excited signals.

The workflow of the whole imaging system is basically the repetition of three steps as shown in Fig. 1d: Step (1) the surface layer of the sample with an optical-sectioning thickness provided by UV surface excitation will be quickly stained by the chemical dyes (DAPI and PI) with ∼150 seconds (Supplementary Fig. 4); Step (2) during imaging mode, the motor-*y* will move rightwards until the objective lens is located above the imaged sample. Then, the surface layer will be 2D raster-scanned by the 2-axis motorized stages and all obtained tiles will be automatically stitched in parallel by our lab-developed programs based on MATLAB (MathWorks Inc.); Step (3) during the sectioning mode, the motor-*y* will move leftwards to its original position, leaving enough space for vibratome sectioning. At the same time, the sample holder will move upwards with the same distance as the slicing thickness to maintain the sample surface always at the focal plane of the objective lens. The above 3 steps will be repeated until the whole organ has been completely imaged. To realize fully automated serial imaging and sectioning, a custom driving system based on a microcontroller unit (MCU) (Mega 2560, Arduino) has been developed to trigger and synchronize all hardware, including the 2-axis motorized stages, vibratome, and color camera. A corresponding control interface based on LabVIEW (National Instruments Corp.) has also been developed to adjust key scanning parameters such as the step size and travel range of the motorized stages.

### Sample preparation

Once mice (C57BL/6) were sacrificed, organs or embryos inside were harvested immediately and rinsed by phosphate-buffered saline (PBS) solution for a minute. Then, the organs or embryos will be submerged under 10% neutral-buffered formalin (NBF) at room temperature for 24 hours for fixation.

To achieve high sectioning quality, it is common to embed the tissue samples into agarose. For different types of tissues, the suggested concentration of the agarose and the corresponding working parameters of the vibratome will be varied^81^. To realize stable performance when sectioning hard organs (e.g., a fixed mouse spleen), 10% (w/v) gelatin is preferred for embedding because post-fixation by NBF overnight can tighten the connection between the sample surface and surrounding gelatin after crosslinking^82–84^. Otherwise, the sample can be pulled out from the embedding medium during sectioning. Moreover, because of the porous structure of lung tissue, perfusing it with low-temperature agarose or gelatin through its trachea is necessary to provide adequate support inside the tissue.

All mice were supplied by the Animal and Plant Care Facility at the Hong Kong University of Science and Technology (HKUST). All the experiments are under the approval of the Animal Ethics Committee and the medical surveillance of the Health, Safety and Environment Office in HKUST.

### Image processing and visualization

The electrical signal from MCU synchronizes the motor scanning and triggers the color camera to capture and save the current FOV as a binary file through its built-in program. A lab-built program based on MATLAB has been developed to further convert binary files to TIFF images and downsize them into different scales (4×, 25×, and 50×) in real time to facilitate the low-resolution 3D rendering later. Also, the program will automatically start to stitch all mosaic FOVs since the current section of the sample has been fully scanned in order to monitor imaging results during the experiment. This feature could highly improve the image processing efficiency, considering the large size of the whole dataset (e.g., ∼2 TB for a whole mouse brain).

Image registration is not necessary for stitching because relative positions between FOVs are well determined by the driving parameters of the high-precision stages. Another reason is that the cross-sections are mostly flat due to the reliable performance of the vibratome. To avoid a grid-like shading pattern on the stitched section due to uneven illumination, conventional flat-field correction^85^ and linear blending algorithm^86^ have been applied. If necessary, notching filters in Fourier-domain can be used to further eliminate the periodic stripes. The color dynamic range of images can also be further adjusted to enhance the contrast. The removal of the background surrounding the sample can be realized by manual masking to achieve high accuracy and minimize artifacts.

To render the 3D model, acquired 2D sections can be directly input to Amira (Thermo Fisher Scientific Inc.) or ImageJ (NIH)^87^ with a *z*-axis interval set as the sectioning thickness. Limited by the computational power, a low-resolution mode is generally applied when rendering the whole block and only a small zoomed-in region will be rendered in high resolution to show more details. 3D structures of some biological components can be extracted based on their colors or signal intensities. For example, nerve fibers in the brain appear to be bright in blue/green channels (Fig. 2t–v), cell nuclei appear to be bright in the red channel (Fig. 2q–s), and white pulps in the spleen appear to be bright in blue/green channel (Fig. 3v). Also, the excited signal from vessels is obviously lower than that from surrounding tissues. Therefore, negative contrast of the vessel network in organs (Fig, 2w,x; Fig. 3j,o) can be utilized for segmentation. All extracted structures have been rendered with pseudo color to enhance their visibility.

### Real-time molecular staining

Typically, a specimen should be washed with PBS after staining to remove residual unbonded probes on its surface, which can induce a strong background signal. However, in TRUST, samples need to be immersed into the staining solutions during the whole experiment for real-time staining. Therefore, the dyes we used should have high specificity, and the intensity of the excited fluorescence signal should only be strong when the probes have been attached to the targets. For example, once bonded to nucleic acids, DAPI or PI exhibits an over 20-fold stronger fluorescence emission than that of unbonded state^47,48^. Since the staining speed of PI is much faster when compared with DAPI^10^, the staining time needed for each round can be simply quantified by analyzing changes in the total number of cells labeled by DAPI over time, which is approximately 150 seconds (Supplementary Fig. 4).

For small samples, the concentration of the staining solutions in the tank will not change significantly throughout the entire imaging process. However, for a large sample, e.g., a whole mouse brain, additional chemical dyes should be added into the water tank by a pump during imaging to maintain the concentration of the solution to be stable. The starting concentration of the staining solutions and the pumping rate applied for different samples are listed in Supplementary Table

1. Because the change of the concentration of solutions is relatively slow, in practice, the control of the pumping rate can be manually tuned based on the average brightness of the imaged sections. In the future, a negative feedback control system can be developed to automatically adjust the pumping rate according to the brightness of the imaged section in real time. It is also possible to simply replace the water tank of the vibratome with one that can hold a larger volume of staining solutions.

## Code availability

The code for detectron2 is available at https://github.com/facebookresearch/detectron2.

## Data availability

The authors declare that all data supporting the results in this study are available from the corresponding author upon request.

## Acknowledgments

The Translational and Advanced Bioimaging Laboratory (TAB-Lab) at HKUST acknowledges the support of the Research Grants Council of the Hong Kong Special Administrative Region (16208620 and 26203619).

## Author contributions

W. Y. and T. T. W. W. conceived of the study. W. Y. built the imaging system. L. K. wrote the control software. W. Y. and V. T. C. T. prepared specimens involved in this study. W. Y. performed imaging experiments. W. Y., Y. Z., and I. H. M. W. performed the image processing. W. Y. and T. T. W. W. wrote the manuscript. T. T. W. W. supervised the whole study.

